# Nanopore sequencing enables high-resolution analysis of resistance determinants and mobile elements in the human gut microbiome

**DOI:** 10.1101/456905

**Authors:** Denis Bertrand, Jim Shaw, Manesh Kalathiappan, Amanda Hui Qi Ng, Senthil Muthiah, Chenhao Li, Mirta Dvornicic, Janja Paliska Soldo, Jia Yu Koh, Ng Oon Tek, Timothy Barkham, Barnaby Young, Kalisvar Marimuthu, Chng Kern Rei, Mile Sikic, Niranjan Nagarajan

## Abstract

The analysis of information rich whole-metagenome datasets acquired from complex microbial communities is often restricted by the fragmented nature of assembly from short-read sequencing. The availability of long-reads from third-generation sequencing technologies (e.g. PacBio or Oxford Nanopore) can help improve assembly quality in principle, but high error rates and low throughput have limited their application in metagenomics. In this work, we describe the first hybrid metagenomic assembler which combines the advantages of short and long-read technologies, providing an order of magnitude improvement in contiguity compared to short read assemblies, and high base-pair level accuracy. The proposed approach (OPERA-MS) integrates a novel assembly-based metagenome clustering technique with an exact scaffolding algorithm that can efficiently assemble repeat rich sequences. Based on evaluations with defined *in vitro* communities and virtual gut microbiomes, we show that it is possible to assemble near complete genomes from metagenomes with as little as 9× long read coverage, thus enabling high quality assembly of lowly abundant species (<1%). Furthermore, OPERA-MS’s fine-grained clustering is able to deconvolute and assemble multiple genomes of the same species in a single sample, allowing us to study the complex dynamics of the human microbiome at the sub-species level. Applying nanopore sequencing to gut metagenomes of patients undergoing antibiotic treatment, we show that long reads can be obtained from stool samples in clinical studies to produce more meaningful metagenomic assemblies (up to 200× improvement over short-read assemblies), including the closed assembly of >80 putative plasmid/phage sequences and a 263kbp jumbo phage. Our results highlight that high-quality hybrid assemblies provide an unprecedented view of the gut resistome in these patients, including strain dynamics and identification of novel plasmid sequences.

## Introduction

The human gut microbiome is known to harbor a rich microbial community composed of hundreds of species with diverse metabolic properties that contribute to multiple facets of host health^1^. In addition, its role as a gene reservoir, particularly in the context of antibiotic resistance genes, has been well documented^2^. The presence of trillions of bacteria cohabiting in close proximity, and under constant selection through diet and occasionally through antibiotics, creates a dynamic environment where antibiotic resistance genes are readily transferred between bacterial species^3^. With an increasing prevalence of multi-drug resistant organisms, many of which carry resistance elements on plasmids transmissible across broad classes of bacteria (e.g. Carbapenem-resistant Enterobacteriaceae: CRE), the role of the gut microbiome as a reservoir that facilitates transmission of antibiotic resistance is of significant scientific and public health interest.

Studies of the transmission of antibiotic resistant organisms between hosts and the environment have primarily relied on cultured isolates, though increasingly metagenomic techniques are being applied to this problem^4–7^. By avoiding culture bias, shotgun metagenomic sequencing promises a more complete view of the gut microbiome, including antibiotic resistant bacteria or fungi and corresponding genes. In practice, however, the limitations of short second-generation reads and the complexity of the metagenome assembly problem, particularly when multiple related species and strains are present in a community, can lead to inaccurate or incomplete assemblies^8^. Clustering methods that use information from sequence composition and coverage provide a valuable approach to aggregate fragmented assemblies into “bins” that likely come from a single species^9,10^.Another major development in this area has been the extension of these ideas to analyze data from multiple samples to improve binning and assemble near complete species-level genomes^11–14^. These methods were tailored to uncover consensus genomes for novel species, but not specifically to assemble different strains and the auxiliary genes in them correctly^15^. As regions of variation across strains can serve as informative markers for transmission studies^16^, there is still a need for methods that can resolve metagenomic assemblies at the strain level in individual samples.

The use of long-read sequencing technologies is a direct way to resolve ambiguity in metagenomic assemblies for individual samples and boost assembly contiguity^17,18^. Its use has however been limited until now due to several conditions including the high cost of sequencing, sequencing biases, stringent DNA requirements and low sequence quality. Recently, ultra-long reads from nanopore sequencing have been used to improve the assembly of a human genome^19^, and significant improvements in throughput have been reported through the introduction of new platforms such as the PromethION. Nanopore sequencing has also been used for metagenomic profiling for some sample types^20^, but its use for metagenomic assembly particularly for human gut microbiome studies has not been explored. In this work, we establish the feasibility and utility of nanopore sequencing for stool metagenomics in clinical studies to obtain long reads that reliably represent the diversity of the metagenome. By combining nanopore and Illumina reads through a first-of-its-kind hybrid metagenomic assembly algorithm, we exploit the strengths of both sequencing approaches and show that we can meet the previously unachievable goals of high base-pair accuracy and near-complete genomes while resolving assemblies at the sub-species level. We demonstrate that these assemblies can then serve as valuable references for studying the evolution of gut resistomes, including strain dynamics and the identification of novel plasmid sequences.

## Results

### High quality long read metagenomics and hybrid assembly with nanopore sequencing

To evaluate the feasibility of routine long read metagenomic sequencing with stool samples from clinical studies, we extracted DNA and assessed its quality in a set of 197 samples from an ongoing study on CRE colonization of the gut microbiome. As is typically the case for such studies, we worked with small quantities (0.5g) of frozen stool samples but noted that despite this limitation, sufficient (>2.5µg) and high-molecular weight (modal size >1kbp) DNA could be obtained for >60% of the samples using our adapted protocol (**Methods**; **Supplementary Figure 1**). In addition, for >25% of the samples we obtained modal DNA size >5kbp, which is well-suited for long read sequencing. A subset of samples (n=28) were then sequenced on a single MinION/GridION flow-cell each (**Supplementary File 1**), generating up to 8Gbp of sequence data per sample, with median throughput >2.5Gbp (**Supplementary Figure 2A**). Read length distributions had a median N50 of 2kbp, and on average >15% of the sequence was in reads longer than 5kbp, making the data suitable for improving contiguity in metagenomic assemblies (**Supplementary Figure 2A** and **Supplementary File 1**). In addition, we noted that the taxonomic distributions for nanopore libraries were highly concordant with those obtained using Illumina short-read libraries (median Pearson correlation >0.95; **Supplementary Figure 2B**) indicating that our nanopore protocols do not induce distinct strong biases for specific bacteria. Overall, our results support the routine use of nanopore sequencing for obtaining long reads to do metagenomic assembly in clinical gut microbiome studies.

To assemble this new type of metagenomics data, we designed the first hybrid metagenomic assembler for long error-prone reads (OPERA-MS), with the aim to obtain highly contiguous assemblies with low base-pair error. OPERA-MS is based on a workflow that is designed to leverage existing tools while addressing the key algorithmic challenge of disentangling related genomes accurately (**Supplementary Figure 3**), using a clustering approach that takes into account assembly graph as well as coverage information to optimize the Bayesian Information Criterion (BIC) of clusters (**Methods**). An overview of the OPERA-MS workflow is shown in **Figure 1** with the following key components: (i) Assembling preliminary contigs using a short-read metagenomic assembler (e.g. Megahit^21^, metaSPAdes^22^, IDBA-UD^23^) and overlaying long-read information to construct an assembly graph of all genomes (steps 1 and 2), (ii) disentangling genomes at the species level using a reference-free clustering approach and augmenting clusters using a reference guided approach (steps 3, 4 and 5), (iii) identifying sub-species level clusters and scaffolding and gap-filling them using OPERA-LG^24^ (steps 6, 7 and 8). We next evaluated OPERA-MS’s ability to leverage long error-prone reads to significantly enhance assembly contiguity while maintaining low assembly error rates compared to state-of-the-art assemblers.

**Figure 1:**
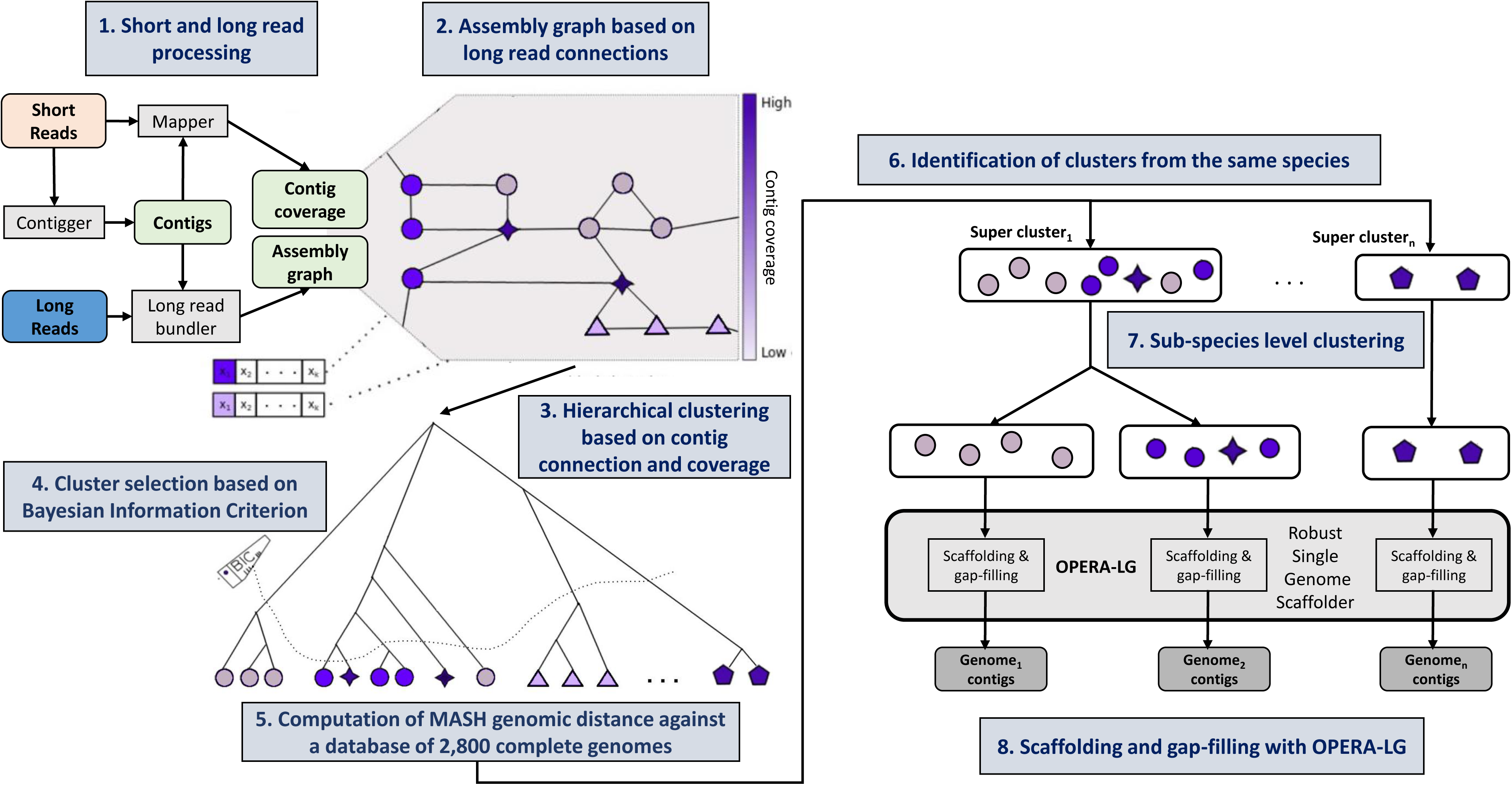
Schematic depicting the steps and workflow for OPERA-MS. Short reads are first assembled by a metagenomic assembler into contigs, and short and long reads are mapped to them to obtain coverage information and spanning reads (Step 1). Spanning reads are then bundled to get edges between contigs for an assembly graph that represents the contiguity information of the whole metagenome (Step 2). Contigs are organized into a hierarchical clustering where the distance between contigs increases with genomic distance and their difference in coverage (Step 3). The tree is then cut into optimal clusters based on the Bayesian Information Criterion (Step 4). To improve the clustering for species where a reference genome is available, the MASH genomic distance between each cluster and a database of complete bacterial genomes is computed (Step 5). Clusters are then merged if there is supporting information in the assembly graph to form species-specific super clusters (Step 6). These super clusters are further analyzed to deconvolute contigs that come from distinguishable sub-species genomes (Step 7). Finally, each cluster is independently scaffolded and gapfilled using a program meant for isolate genomes (OPERA-LG; Step 8).

### Recovering near-complete and high fidelity genomes from hybrid metagenomic datasets

To comparatively evaluate the performance of various programs, we assembled sequencing datasets for *mock communities* where the ground truth for the metagenome is well known. This included the HMP staggered mock community for which Illumin short reads^18^ a, Illumina Synthetic Long Reads^18^ and PacBio data are publicly available, as well as a new diverse community of 20 species (GIS20) whose abundances ranged from 0.1 to 30% and were deeply sequenced on the Illumina (80% species with >30× read coverage), PacBio and Oxford Nanopore (55% and 65% species with read coverage >5×, respectively) platforms (**Supplementary Table 1**, **Supplementary File 2** and **Supplementary Figure 4**; **Methods**). For both communities we noted that the data adequately captured community composition, with high correlation between the expected and observed relative abundance of species for Illumina and Oxford Nanopore data (Spearman ρ>0.96), slightly lower correlation for PacBio data (0.94-0.97), and lower but still high correlation for Illumina Synthetic Long Reads (0.89; **Supplementary Figure 4**). Overall, we evaluated three state-of-the-art metagenomic assemblers (MegaHit^21^, metaSPAdes^22^, IDBA-UD^23^), a long-read assembler (Canu^25^, not specifically designed for but used for metagenomic assembly^25^), a hybrid assembler (hybridSPAdes^26^, with test extensions for metagenomics) and the hybrid metagenomic assembler OPERA-MS, on metagenomic data representing 37 completely assembled bacterial genomes across 4 different sequencing approaches (Illumina, PacBio, Oxford Nanopore and Illumina Synthetic Long Read).

We first began by assessing the relative benefits of short and long read sequencing for metagenomic assembly based on these gold-standard datasets. Currently, short-read Illumina sequencing is still the most cost-effective approach to study complex microbial communities. We noted that as expected, the ability to sequence deeply does improve metagenomic assembly, but assembly contiguity (measured by NGA50 per genome, with assembly errors accounted for; **Methods**) plateaus out at ~30× coverage (NGA50 <300kbp with metaSPAdes; <200kbp for MegaHit and IDBA-UD; **Figure 2A**, **Supplementary Figure 5**). This limitation in assembly contiguity is however addressed by longer reads that can span repetitive sequences in microbial genomes, and correspondingly near complete genomes (NGA50 >1Mbp) are obtained even from metagenomic data when sufficient coverage is available (ideally >60×; **Figure 2B**). The precise requirements for long read coverage vary across genomes due to the specifics of their repeat content, but at a minimum, assembly with Canu required >10× coverage and many genomes remained unassembled even at this coverage (**Figure 2B**). In contrast, assembly with hybrid approaches that leveraged both short and long read data enabled NGA50 >100kbp with as little as 5× long read coverage (**Figure 2C** and **Supplementary Figure 5**). Notably, OPERA-MS assembled near complete genomes with ~9× long read coverage while Canu and hybridSPAdes required >30× coverage to obtain similar quality assemblies. In addition, base-pair level accuracy was significantly lower for metagenomes assembled solely with native long reads (Canu), primarily due to 10-100× increase in median indel error rate, even after consensus correction (**Methods**; **Supplementary Figure 6**). Hybrid assembly methods (OPERA-MS, hybridSPAdes) provided higher sequence quality comparable to some short read methods (median quality ≈ Q40), further highlighting the advantages they provide over short read and long read only approaches for metagenome assembly contiguity and sequence quality, respectively.

**Figure 2:**
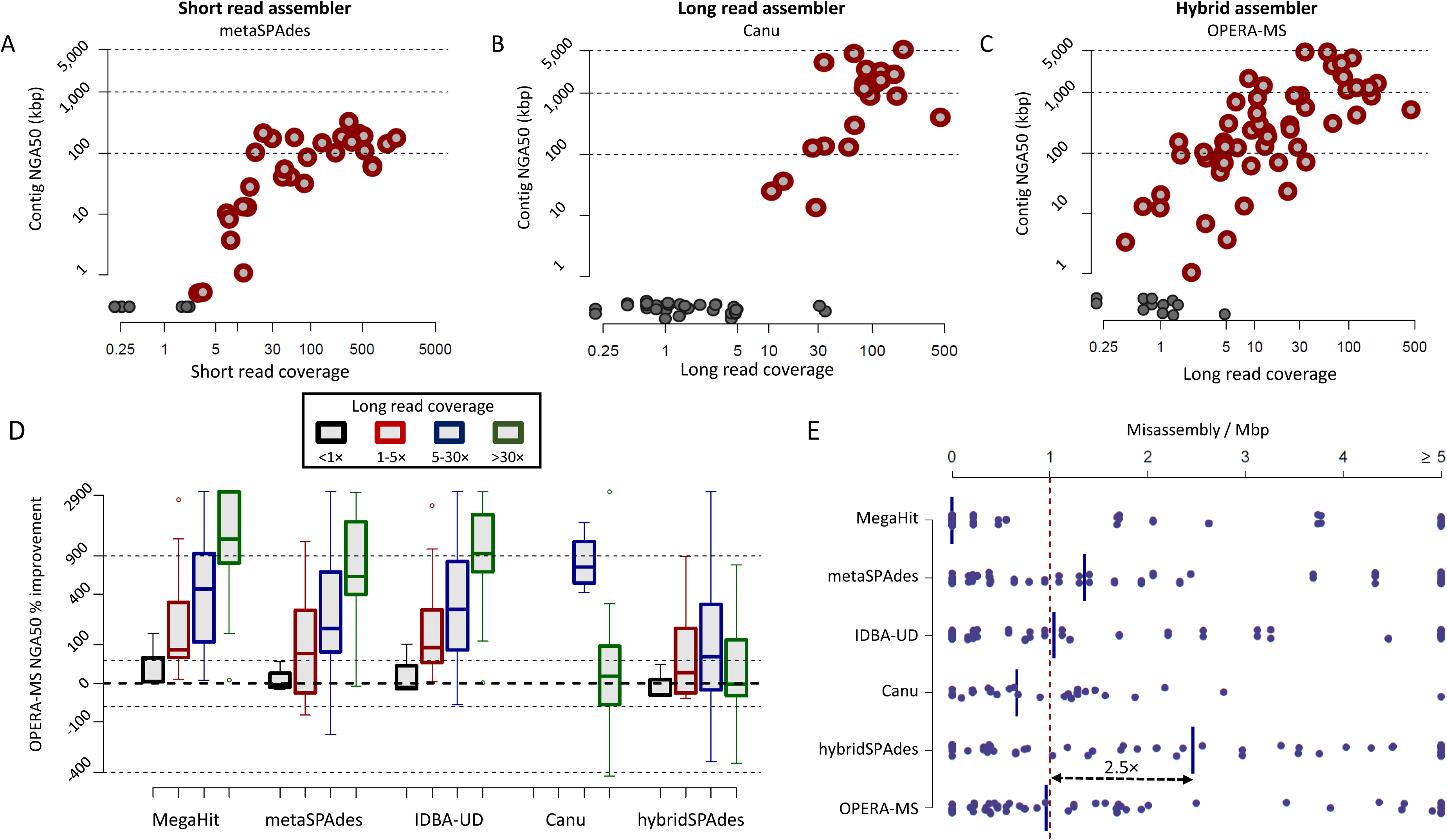
Hybrid assembly of near-complete and high-quality genomes from metagenomic datasets. Increase in assembly contiguity as a function of read coverage for a representative, **A** short read assembler, **B** long read assembler, and **C** hybrid assembler. Note that hybrid assembly improves over short and long read assembly in terms of scaling across coverage ranges and producing near-complete genomes (NGA50 >1Mbp) with as little as 9× long read coverage. Unassembled genomes are shown as circles with black borders. **D** Improvements in assembly contiguity (NGA50) provided by OPERA-MS in comparison to other assemblers as a function of long read coverage. Note that Canu does not assemble low coverage genomes and hence metrics are not provided in those ranges. **E** Misassembly rates for different assemblers. Most assemblers produce ~1 large misassembly per Mbp, except for hybridSPAdes.

To analyze OPERA-MS’s utility as a function of long read coverage in comparison to other assemblers, relative improvements in assembly contiguity (NGA50) were computed per genome for different coverage bins (<1×, 1-5×, 5-30×, >30×). Compared to short-read assemblers, OPERA-MS enables notably better assembly contiguity (>50%) as long as coverage is >1× (~10- fold improvement with coverage >30×; **Figure 2D**), and these improvements were seen with assemblies from metaSPAdes and IDBA-UD as well, in addition to MegaHit (**Supplementary Figure 7**). In relation to long read only assemblies (Canu), OPERA-MS assembled many highly contiguous genomes (NGA50 >100kbp) with <5x long read coverage where Canu is unable to assemble genomes (**Figure 2B**, **C**), provided >800% improvement for genomes with 5-30x coverage, and similar contiguity at higher coverage (>30×). Among hybrid approaches, OPERA-MS improved over hybridSPAdes assemblies by >50% for 40% of the genomes assembled, compared to <5% of genomes with such improvements using hybridSPAdes versus OPERA-MS. Of note, median improvement in assembly contiguity was more than 100% for genomes with 5- 30× coverage when comparing OPERA-MS with hybridSPAdes. The advantages of hybrid assembly were also seen in terms of assembly completeness with OPERA-MS assembling an additional 20-40kbp of sequence on average per genome compared to short read assemblies (**Supplementary Figure 8**). Finally, we noted that OPERA-MS typically produces <1 misassembly per Mbp, a 2.5× improvement over hybridSPAdes which produced the most errors among the tested methods as expected for an isolate genome assembler^18^ (**Figure 2E**). Overall, these results indicate that OPERA-MS is a versatile approach capable of obtaining highly contiguous and accurate genome assemblies from diverse long read metagenomic datasets.

### OPERA-MS accurately assembles strain genomes in complex communities

While mock communities are a commonly used gold-standard for evaluating metagenomic assembly with real reads, they typically represent simpler communities. To address this and evaluate assembly methods on more complex communities where ground truth is known we took the approach of spiking mock community reads (GIS20) into our stool metagenomics data (sample S2, **Supplementary File 1**) to construct *virtual gut microbiomes* (**Figure 3A**). These virtual gut microbiomes carry the complexity of real datasets while allowing us to evaluate OPERA-MS’s accuracy in assembling strain genomes when multiple strains are present in the data. In particular, we noted that GIS20 shared five species with the stool dataset that it was spiked into (*Klebsiella pneumoniae*, *Streptococcus parasanguinis*, *Enterobacter clocae, Bifidobacterium adolescentis* and *Parascardovia denticolens*) and these would be expected to have their assembly impacted in the virtual community.

**Figure 3:**
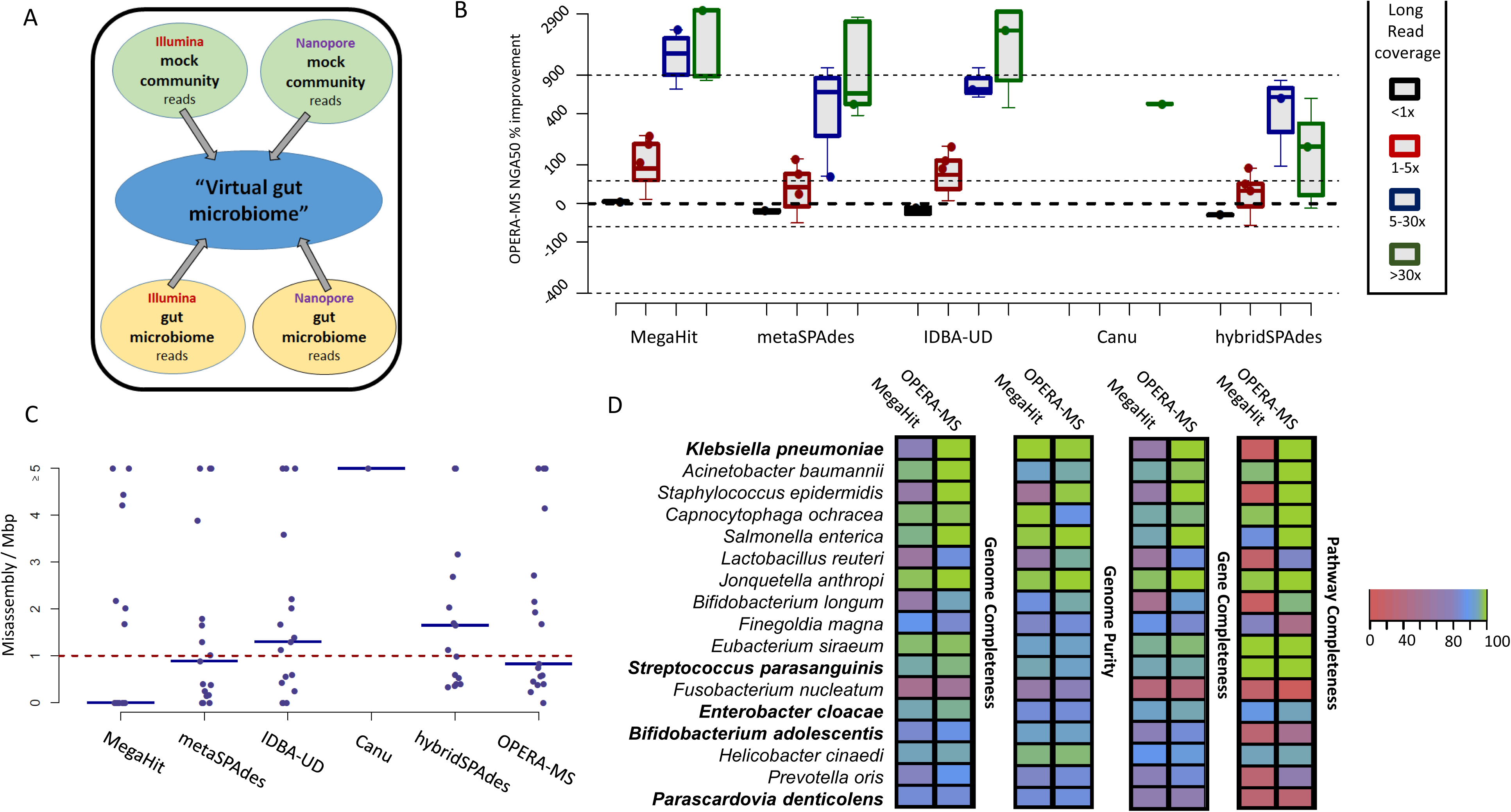
OPERA-MS improves assembly and analysis of a complex metagenome from a virtual gut microbiome. A. Construction of a virtual gut microbiome that represents a complex metagenomic dataset while retaining the ability to evaluate assemblies against gold-standard references. **B** Improvement in assembly contiguity (NGA50) obtained using OPERA-MS compared to other assemblers over different coverage ranges. Dots represent species that have at least two strains in the metagenome (i.e. present in GIS20 and S2 with an abundance >0.1% as reported by MetaPhlAn2^72^ (v2.6.0)). **C** Comparison of misassembly rates for different assemblers. **D** Evaluation of Illumina-only (MegaHit) and hybrid (OPERA-MS) metagenomic assemblies after binning for their utility in downstream analysis. Bins that contained the largest fraction of a reference genome (GIS20 references; species with bold names have at least two strains in the metagenome) were evaluated for (i) genome completeness = the fraction of the genome represented in the bin, (ii) genome purity = percentage of bases in the bin that correspond to the correct reference, (iii) gene completeness = fraction of genes that were fully assembled in the bin, and (iv) pathway completeness = fraction of pathways with over 90% of their constituent genes being assembled and binned together.

Even with the increased diversity of the community (Shannon divergence 3.4 *vs* 2.1 for GIS20), OPERA-MS continued to exhibit 5-10 fold improvement in assembly contiguity compared to short read metagenomic assemblers (MegaHit, metaSPAdes, IDBA-UD) with genome coverage >5× and a useful 50% improvement with 1-5× coverage (**Figure 3B**). The utility of metagenomic assembly methods was also more apparent in the virtual gut versus the mock community datasets. While Canu assembled only one genome and at 5-fold lower contiguity than OPERA-MS, contiguity was 2-fold higher with >5× coverage for OPERA-MS *vs* hybridSPAdes assemblies. These differences were also apparent in terms of misassembly, with Canu and hybridSPAdes having >1.5× of the misassembly rate obtained with OPERA-MS (**Figure 3C**).

As “binning” of fragmented short read metagenomic assemblies is commonly performed to aggregate sequences that may come from the same genome for downstream analysis, we next evaluated results with this step. Applying MaxBin2^10^ to the most accurate short-read (MegaHit) and long-read (OPERA-MS) assemblies revealed substantial difference in binning quality (**Figure 3D**). Median genome completeness with OPERA-MS was improved to 95% *vs* 83% with MegaHit (median genome purity of 90% for both methods), with four (out of 20) OPERA-MS bins with genome completeness >99% and purity >95% (0 for MegaHit). Binning performance also markedly impacted analysis of (i) *gene content*, with 93% gene assembly with long reads and OPERA-MS *vs* 82% with short reads, as well as (ii) *pathway reconstruction*, with 96% coverage with OPERA-MS *vs* 61% with MegaHit (**Figure 3D**; **Methods**), highlighting the value of hybrid metagenomic assemblies for gene content and metabolic pathway analysis^27^.

As a specific example of assembly and binning performance differences, we observed that despite having high short read coverage (>100×) in the virtual gut microbiome, MegaHit assembly of the *Klebsiella pneumoniae* strain that was spiked in from GIS20 (10× more abundant than native strains) had NGA50 of ~11kbp, a 5-fold reduction compared to its GIS20 assembly. Binning of the assembly led to three different bins, each containing a fraction of the genome (50%, 25% and 15%). This is concordant with known challenges for metagenomic assembly contiguity and accuracy in the presence of multiple strains of the same species^8^. Even when manual binning was performed to combine the 3 bins, the Illumina-only *K. pneumoniae* assembly still did not have ~20% of its genes assembled, including 2 out of the 3 antibiotic resistance genes that are known to be present in this strain (**Supplementary Figure 9A**). Assembly with Canu was able to leverage the >100× long read coverage to produce 12 contigs covering ~93% of the genome, but with 53 relocation errors representing 419kbp of misassigned novel sequence (NGA50 = 152kbp). Similarly, the hybridSPAdes assembly had NGA50 of 327kbp and 448kbp of novel sequence in 10 relocation errors. The OPERA-MS assembly generated a single 5Mbp contig with 5 relocation errors and 6kbp of novel sequence (NGA50 = 1.4Mbp), representing a >120× improvement in contiguity over the initial MegaHit assembly. In total, 99% of the genes were fully assembled by OPERA-MS, including all antibiotic resistance genes, and the genome was contained in a single high quality bin (99.8% completeness and 99.3% purity) which almost fully captures the pathways of this *Klebsiella pneumoniae* strain (**Supplementary Figure 9B**). These results showcase the utility of hybrid metagenomic assembly for gene content and pathway analysis in complex microbial communities and the ability to accurately distinguish sub-species genomes in them with OPERA-MS.

### Hybrid assembly for analysis of antibiotic resistance and novel mobile elements in the human gut microbiome

Based on the evidence that OPERA-MS can accurately resolve strain ambiguity and assemble complex metagenomes, we next applied it to nanopore and Illumina sequencing data for stool samples from our initial cohort (n=28; **Supplementary File 1**, **Supplementary Figure 2**; **Methods**). Samples for this cohort came from two studies that investigate the impact of antibiotics^28^ and CRE colonization (manuscript in preparation) on the gut microbiome, and correspondingly we were interested in additional insights into mobile elements and antibiotic resistance that we could obtain from hybrid metagenomic assembly. In terms of overall assembly, while our initial MegaHit assemblies had median N50 <9kbp, hybrid assembly resulted in >5× improvement in median assembly contiguity, with an even larger improvement for lowly abundant species (10×, starting from 4kbp), and up to 200× improvement for some genomes (**Supplementary Figure 10**). In terms of high quality genomes as assessed by CheckM^29^, OPERA-MS nearly doubled the number with N50 >100kbp compared to hybridSPAdes (69 *versus* 36), and was able to assemble many high quality and contiguity genomes even in the presence of strain ambiguity (8 *versus* 1 and 0 for hybridSPAdes and MegaHit, respectively; **Supplementary Figure 11**; **Methods**). The average runtime on these datasets was <12 hours for the combination of MegaHit and OPERA-MS using 20 CPUs, in comparison to slightly more than 12 hours for hybridSPAdes.

Due to the presence of repetitive elements, assembly of mobile elements with short reads can be challenging even in isolate genome data^30^. Using circularity of assembled sequences to identify potential fully assembled phage and plasmid genomes, we directly recovered 8.9Mbp of circular sequences in 88 contigs from OPERA-MS gut metagenome assemblies (**Methods**). The assembled sequences covered a wide range of sizes (16 sequences > 100kb; **Figure 4A**), including a complete bacterial genome, large plasmids and phages (**Supplementary File 2**), none of which were fully assembled using only short reads (**Supplementary Figure 12A**). The assembly of a complete genome for an *Enterobacter cloacae* strain exhibited high sequence (identity >99%) and structural (no inversions or translocations observed) similarity with the reference genome for *Enterobacter cloacae ENHKU01* (**Supplementary Figure 12B**), indicating high fidelity for these assemblies. Among assembled sequences, a majority (68/88) are substantially diverged from known sequences (average identity <75% or fraction of sequence covered <85%; **Figure 4B** and **Supplementary File 2**) while 18 sequences appear to be completely novel (no blast hit against nt database), highlighting the potential to uncover novel plasmids and phages in the human gut microbiome through hybrid metagenomic assembly^31^.

**Figure 4:**
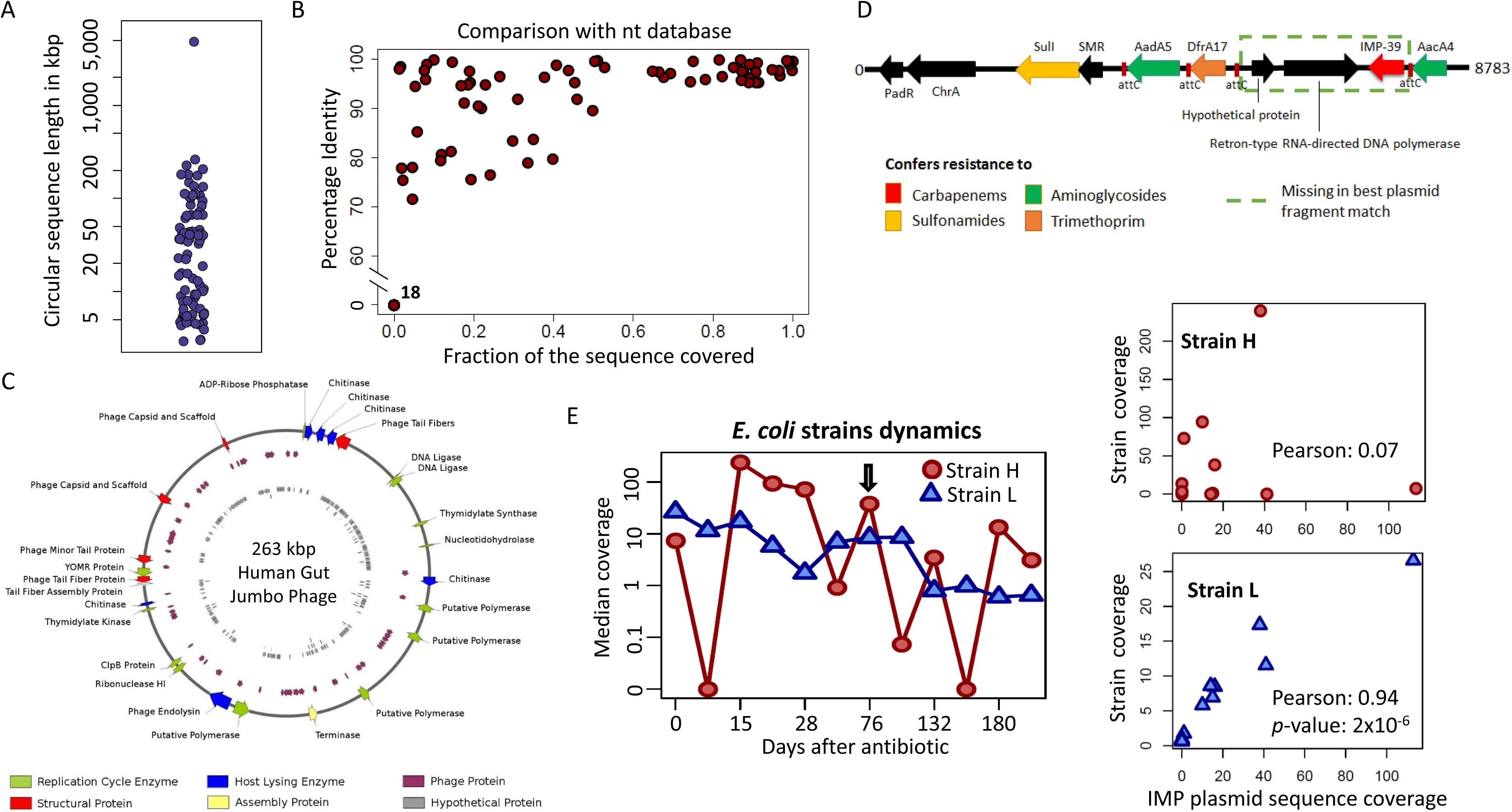
Assembly of novel mobile elements and their association with host species in the human gut microbiome. **A** Distribution of genomes sizes for fully assembled circular sequences from OPERA-MS in 28 human gut metagenome datasets, illustrating the ability to assemble circular genomes of varying sizes and complexity (plasmids, phages and bacterial genomes). **B** Fraction of sequence covered *versus* average sequence identity of the assembled circular sequences in comparison to sequences in the nt database (based on BLAST searches). Many of the assembled sequences showed good alignment and homology to known sequences from end to end (top right corner), but some only had local similarities (top left corner), and a few appear to be completely novel (bottom left corner; 18 sequences). **C** Annotation of the largest (263kbp) observed novel circular sequence (no matches in nt database) revealed proteins associated with a phage life cycle, including replication, assembly and host lysis, indicating that the assembled sequence is a putative jumbo phage. **D** A novel multiple resistance region assembled by OPERA-MS from the gut microbiome of a CRE-infected patient. Apart from the clinically relevant carbapenamase gene cassette, the region also harbors genes that confer resistance to aminoglycosides, trimethoprim and sulfonamides, limiting treatment options. **E** Strain level assembly with OPERA-MS enabled association of plasmid to genome based on correlation in read coverage across timepoints. Left panel: Variation in coverage of 2 *E. coli* strain genomes seen in the hybrid metagenomic assembly of data from day 76 (black arrow). Right panel: Correlation between the coverage of the plasmid and the 2 *E. coli* strains reveals that it is strain L that likely harbors the IMP gene containing plasmid.

Further characterization of the largest novel circular sequence (263kbp) confirmed that the assembly is well support by read data, with uniform coverage across the entire sequence of long ONT reads (>1kbp) that align end-to-end (**Supplementary Figure 13A**). Annotation of the genome revealed the presence of multiple phage genes (>50), including those related to the phage tail and tail assembly protein, capsid protein, polymerase, chitinase, terminase and ligase, all of which are essential proteins for the life cycle of a phage (**Figure 4C**). In addition, no plasmid related genes were found, providing evidence for this sequence representing the first genome of a jumbo phage^32^ (genome size > 200kbp) to be identified in the human gut microbiome. Phylogenetic analysis further supported this hypothesis, with the polymerase gene in this genome grouping with other jumbo phage genes in a cluster that was well separated from polymerase genes for small phages and bacteria (**Supplementary Figure 13B**). Interestingly, the putative jumbo phage was abundant in the gut metagenome of an individual while receiving antibiotic treatment (and at no other timepoints preceding or following that; **Supplementary Figure 13C**), and was accompanied by an Enterobacteriaceae (*K. pneumonia, E. coli and C. freundii*) bloom. Two other large putative coliphages were also identified in our assemblies, a 150kbp sequence with 96% identity to *phAPEC8* over 89% of its genome and a 93kbp sequence with 96% identity to *P1 mod749* over 82% of its genome, showcasing the promise of mining gut metagenomes for phages that may be useful in the fight against antibiotic resistant bacteria^33^.

In order to study antibiotic resistance (AR) gene combinations, we expanded our analysis to non-circular sequences and annotated >1.68 Gbp of contigs longer than 5kbp (**Methods**). In particular, as many samples in the study are CRE colonized and treatment options for CRE infections are limited^34^, the direct detection of specific resistance combinations was of interest to us to understand its utility for guiding therapy. Leveraging on the sequence contiguity provided by our hybrid assemblies, we discovered 20 (out of 130) novel combinations of AR genes (ranging from 2 to 7 genes) that are not represented in current public genome databases (**Supplementary Table 2**), and 4 of which harbor a carbapenamase (2 Class A, 1 Class B and 1 Class D). We noted that several carbapenemase genes despite likely being capable of hydrolysing most β-lactams^35^, still co-occurred with extended spectrum β-lactamases (**Supplementary Table 2**), corroborating recent reports on the increasing incidence of bacterial isolates with multiple β-lactamase genes^36^. These co-occurences could contribute to increased resistance to β-lactams^37^, potentially impacting combination therapies for CRE infections involving β-lactams^38^ or new β-lactam antimicrobials (e.g. Ceftazidime/avibactam).

Non-β-lactam antibiotics such as aminoglycosides and trimethoprim-sulfamethoxazole are increasingly considered for use in carbapenem-resistant gram-negative bacterial infections^39^. Of note, we assembled a multiple resistance region (MRR) that harbors a previously unknown combination of AR genes for carbapenems, aminoglycosides, trimethoprim and sulfonamides (**Figure 4D**, **Supplementary Table 2**). The MRR contains multiple repeats that fragment its Illumina assembly, while alignment of the hybrid assembly against the best plasmid match suggests that this combination arose by bringing together a gene cassette (demarcated by attC sequences) containing the carbapenamase gene IMP-39 (**Figure 4D**) with a known class 1 integron harboring the chrA and padR genes^40^.

As the MRR containing plasmid was assembled from metagenomic data for a CRE colonized subject and best aligned to a plasmid associated with *E. coli*, we hypothesized that the plasmid was likely hosted by an *E. coli* strain. Among Enterobacteriaceae species, *E. coli* and *K. pneumoniae* were readily detected across 11 timepoints using metagenomic profiling approaches, but their abundances over time were poorly correlated with the abundance of the plasmid (Pearson correlation <0.05; **Supplementary Figure 14**). OPERA-MS assembly, however, disambiguated two *E. coli* genomes that we denoted as strain *H* (high abundance at timepoint for hybrid assembly) and strain *L* (**Figure 4E**). These two strains exhibited very different correlation patterns with the abundance of the IMP plasmid, identifying strain L as the likely host (Pearson correlation 0.94 *versus* 0.07 for strain H; **Figure 4F**). By isolating the CRE strain from stool samples for this subject and sequencing its genome we were also able to confirm the correctness of this association (**Methods**), suggesting that this approach could complement existing Hi-C sequencing based techniques to link plasmids and their host genomes in metagenomic data^41^.

## Discussion

As the cost of long read sequencing continues to go down, its value for metagenomic assembly will become even more compelling. A major concern that has impacted adoption is the ability to work with limited DNA amounts and still generate sufficient number of long reads. Our work establishes that it is indeed feasible to do this with realistic clinical samples on current nanopore systems, using easy to adopt DNA extraction and library preparation protocols. At current throughputs of >10Gbp per MinION flowcell, users can expect >10× long read coverage of even lowly abundant genomes (<1%), allowing them to generate high quality assemblies with OPERA-MS. This can be achieved for around $500 per sample, and multiplexing can help bring these costs down further by leveraging increased throughputs on the GridION/PromethION systems.

Our results highlight that the utility of hybrid metagenomic assembly is similar to what has been noted for isolate genome assembly^42,43^. However, unlike the isolate genome case, challenges in metagenomic assembly are not fully resolved solely with long reads, as their error rates preclude the resolution of similar species and strains in the metagenome, and hamper assembly quality and utility for downstream applications. By drawing on the strengths of both short and long read data, hybrid assembly is the best strategy in such cases, and methods such as OPERA-MS and hybridSPAdes provide a significant advantage over non-hybrid approaches, even for simple communities. For more complex communities such as the ones seen in the human gut, the advantage of a metagenome-specific assembler should be more apparent, as was noted in our results for virtual and real gut microbiomes. The development of OPERA-MS as a first hybrid metagenomic assembler, is thus a useful step in exploiting long read sequencing for “high-resolution” analysis (near-complete genomes and strain resolution) of diverse microbiomes.

In particular, the ability to increase assembly N50s from 10s of kbp to 100s of kbp and even >1Mbp opens up the ability to do new kinds of analysis with metagenomic data. While the metabolic capability of metagenomes is generally analyzed based on a “bag of genes” approach^44^, improved assembly and binning algorithms allow us to correctly account for the outputs of individual genomes and the metabolic interactions that they support^45^. Furthermore, sample specific assembly of the non-core genome, including repeats, transposable elements and plasmids, allows us to better study their evolution and transmission, a capability that is key for tracking antibiotic resistance in the gut microbiome.

In addition to improved contiguity, OPERA-MS assemblies were able to resolve sub-species genomes in a metagenome. This feature allowed us to discriminate between a lowly abundant carbapenem resistant *E. coli* strain and a more abundant sensitive strain in a patient’s gut metagenome - information that can potentially be valuable in a clinical setting. The limitations to resolving sub-species genomes with OPERA-MS are defined by Illumina sequencing error rates and read coverage. Genomic sequences that are either not assembled or collapsed in an Illumina assembly will remain so in OPERA-MS’s output currently, though an obvious extension is to consider haplotype analysis^18^ based on the assembly backbone created by OPERA-MS. In contrast to methods that cluster strain-specific polymorphisms^46,47^, the output from OPERA-MS is tailored to capture structural differences that separate the corresponding strain genomes and thus these methods have complementary utility.

Despite being heavily studied, the human gut microbiome still contains elements that are not well explored such as its “phage reservoir” which is believed to play important roles in its resilience to perturbations^48^, as well as the gut resistome. Based on the hybrid assembly of 28 gut metagenomes, we uncovered >30Mbp of plasmid sequences longer than 20kbp, many of which harbor novel antibiotic resistance cassettes, and some of which have no homology with known sequences. In addition, OPERA-MS assembled several large circular phage genomes, including what appears to be the complete genome of the first putative jumbo phage identified in the human gut microbiome. These results likely represent the “tip of an iceberg” and we envisage that extensive use of hybrid metagenomic assembly will make it increasingly feasible to study resistance determinants and mobile elements in the human gut microbiome.

Clinical metagenomics is an emerging field in which the microbiological diagnosis of complex, polymicrobial infections is determined from sequencing of DNA extracted directly from clinical specimens^6,5,7^. The ability to reassemble plasmids and detect combinations of antimicrobial resistance elements is an important step for molecular surveillance of high impact antimicrobial resistance genes such as carbapenemases as they disseminate and recombine between mobile genetic elements and bacterial species, and for rational antibiotic selection in this treatment paradigm. The enhanced resolution of OPERA-MS assembly thus has the potential to accelerate the implementation of metagenomics into outbreak investigations and surveillance programs where culture-independent approaches speed-up detection of unknown pathogens, novel antimicrobial resistance combinations and virulence mechanisms^6,5,7^.

## Methods

### High molecular weight DNA extraction for stool metagenomics

Metagenomic DNA extraction was carried out using Phenol-Chloroform extraction and various commercial kits (i.e. QIAamp DNA Stool Mini Kit, PowerSoil® DNA Isolation Kit, and PowerFecal® DNA Isolation Kit). The quality and yield of DNA extracted for each technique were compared to assess which method was most suitable. Ease of handling was also taken into account in deciding which method to use.

Phenol-chloroform extraction provided high yields of DNA. However, the integrity of DNA was poor and the process of extraction was more challenging and hazardous compared to commercial kits. QIAamp DNA Stool Mini Kit, PowerSoil^®^ DNA Isolation Kit and PowerFecal® DNA Isolation Kit all generated comparable yields. Although QIAamp DNA Stool Mini Kit was able to provide higher quality DNA, the absence of a bead-beating step in the kit might limit comprehensive lysis for an unbiased metagenomic profile. PowerSoil and PowerFecal DNA Isolation Kits are comparable in all aspects, except for an additional step of heating for lysis and the addition of bead solution in the latter.

To avoid clogging problems and get higher molecular weight DNA for nanopore sequencing, we further optimized the PowerSoil^®^ DNA Isolation Kit protocol. Briefly, samples (250mg of sample per PowerBead tube) were mixed manually by gently inverting the tube several times. Solution C1 was added to tubes and was mixed by inversion. Mechanical lysis was carried out by attaching PowerBead tubes to a vortex adapter and samples were vortexed for 10min at a low speed of 5 (instead of manufacturer’s recommendation of speed 10) to reduce DNA fragmentation. To avoid clogging of spin filters, centrifugation time was extended to twice the original duration and solutions C2, C3 and C4 were doubled in volume. DNA was eluted in 55µL of heated (65°C) solution C6 and concentrated using 1X Agencourt AMPure XP beads (A63882, Beckman Coulter). Purified DNA was quantified by Qubit dsDNA BR assay (Q32853, Thermo Fisher Scientific). DNA size was assessed by Agilent Bioanalyzer (Agilent Technologies) prepared with Agilent DNA12000 Kit (5067-1508, Agilent Technologies). For nanopore sequencing, size selection was applied to all samples with peak size <8kb using 0.45X of Agencourt AMPure XP beads, except sample S10 for which BluePippinTM (Sage science) size selection was completed using a 0.75% dye-free cassette (BLF7503, Sage science) with S1 marker (fragment size between 3–50kbp) followed by cleanup with 1X Agencourt AMPure XP beads. We tested whole genome amplification using the Genomiphi HY Kit (G13/25-6600-20, GE Healthcare) on sample S3 (DNA cleanup with 0.45X Agencourt AMPure XP beads and integrity check on 0.5% agarose gel). However the throughput for this sample was significantly lower compared to other samples (S3, **Supplementary Figure 2**).

### Long read metagenomic sequencing on the MinION

As nanopore sequencing technology advances, various kits, flowcell types and devices were used to sequence the samples as detailed in **Supplementary Table 1**. All libraries were prepared according to the manufacturer’s protocols with minor modifications. All mixing steps for DNA samples were done by gently flicking the microfuge tube instead of pipetting and the optional shearing step was omitted. For all elution steps, an additional microlitre of nuclease-free water or elution buffer was used to avoid any carry over of magnetic beads. DNA repair treatment was carried out using NEBNext FFPE DNA Repair Mix (M6630, New England Biolabs). End-repair and A-tailing was performed with NEBNext Ultra II End-Repair/dA-tailing Module (E7546, New England Biolabs) and samples were incubated at 20°C for 5min and 65°C for 5min. End-repaired product was cleaned up with 1× Agencourt AMPure XP beads (A63882, Beckman Coulter). Adapters provided in the respective library kits were ligated to the DNA with NEB Blunt/TA Ligase Master Mix (M0367, New England Biolabs) and samples were incubated at room temperature for 10min. Purification and loading of adapted libraries was completed as stated in the manufacturer’s protocol and sequenced using the appropriate MinKNOW™ workflow. Libraries were basecalled using Metrichor, Albacore or Guppy (**Supplementary Table 1**) and fastq files were generated using Albacore (samples S6 to S13) or poretools^49^ (all other samples).

### Illumina sequencing of metagenomic libraries

To construct the library, 50ng of DNA was re-suspended in a total volume of 50µL and was sheared using Adaptive Focused Acoustics™ (Covaris) with the following parameters; Duty Factor: 30%, Peak Incident Power (PIP): 450, 200 cycles per burst, Treatment Time: 240s. Sheared DNA was cleaned up with 1.5× Agencourt AMPure XP beads (A63882, Beckman Coulter). Gene Read DNA Library I Core Kit (180434, Qiagen) was used for end-repair, A-addition and adapter ligation. Custom barcode adapters were used in place of Gene Read Adapter I Set for library preparation (HPLC purified, double stranded, 1^st^ strand: 5’ P-GATCGGAAGAGCACACGTCT; 2^nd^ strand: 5’ ACACTCTTTCCCTACACGACGCTCTTCCGATCT). Libraries were cleaned up twice using 1.5× Agencourt AMPure XP beads (A63882, Beckman Coulter). Enrichment was carried out with indexed primers according to an adapted protocol from Multiplexing Sample Preparation Oligonucleotide kit (Illumina). Molarity of libraries were measured using Agilent Bioanalyzer (Agilent Technologies), prepared with Agilent DNA1000 Kit (5067-1504, Agilent Technologies). Enriched libraries were pooled in equimolar concentrations and sequenced on the Illumina HiSeq sequencing platform to generate 2×101bp reads.

### Hybrid metagenomic assembly with OPERA-MS

An overview of the hybrid metagenomic assembly workflow with OPERA-MS can be found in **Figure 1**. In step 1, the short-read data is used to construct contigs, which along with the long read data serves as the input for assembly graph construction. This approach leverages the deeper coverage and higher accuracy of short read sequencing to generate short but accurate contigs, with long reads providing contiguity improvements even with lower read coverage (2-10×) for rare members of complex microbial communities.

#### Assembly graph based on long read connections

Long reads were mapped to the short-read contigs using BLASR^50^ and reads that provide connecting information between contigs were clustered into edges using the procedure in OPERA-LG^24^ with metagenomics specific steps as described below. Recognizing that edges in the assembly graph are more likely to provide incorrect/conflicting information, (i) for metagenomic data due to the presence of multiple closely related species and strain genomes, and (ii) for long read data with higher sequencing/mapping errors and chimeric reads, we tried to identify and resolve such issues. In particular, edge conflicts were identified as contig pairs with multiple edges connecting them that indicate different orientations and distances and only edges with higher support were retained. In addition, edges that had unusually low support were identified by computing the long-read to short-read contig coverage ratio for each edge (*r_e_*) and flagging outliers for removal (< median*_e_*(log(*r_e_*)) − 1.5×std*_e_*(log(*r_e_*))) among edges in different distance classes ([-100, 300], [300, 1000], [1000, 2000], [2000, 5000], [5000, 15000], and [15000, 40000]). Finally, only edges with long read support greater than 1 were kept.

#### Hierarchical clustering based on contig connection and coverage

After assembly graph construction, OPERA-MS applies the strategy of decomposing the metagenomic assembly problem into an isolate genome assembly problem, to both accelerate assembly and leverage the development of robust algorithms for the isolate assembly problem^51^. While existing methods primarily approach this decomposition as a clustering problem based on the read coverage of contigs^52^, the availability of the assembly graph presents a problem akin to image segmentation where we wish to take into account the proximity structure defined by the graph in clustering contigs. This is the core idea in OPERA-MS and is accomplished in two steps using a Bayesian approach: (i) by constructing a “guide tree” along graph edges that connects contigs that have similar coverage and encodes prior beliefs on reasonable clusterings, and (ii) computing an optimal partition of the tree into clusters based on the Bayesian Information Criterion, as detailed below.

We begin with some definitions for the problem that we are trying to solve using a Bayesian approach. Let *X* = {*x*_1_ … *x*_n_} represent the set of contigs assembled from a metagenomic shotgun sequencing project, and *Y* = {*y*_1_ … *y*_n_} represent their corresponding read counts. Then if there are *K* (unknown) components ({*M*_1_ … *M_K_*}, i.e. the corresponding isolate genomes) that underlie the observed count distribution, the probability of observing *Y*, under an independent and identical distributions assumption, is given by a mixture model *M* as

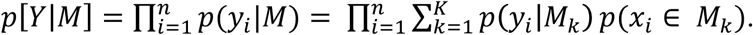

Modelling *p*(*y_i_*|*M_k_*), the number of reads that originate from an isolate genome, is often done with the Poisson distribution. However, the assumption of equality of mean and variance means that this model only works when reads are uniformly distributed on the assembled contigs. Sequencing biases and the presence of repetitive sequences can give rise to over-dispersion and this is typically accounted for by using the negative binomial distribution, where

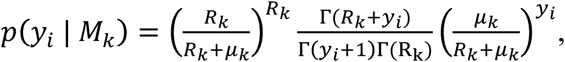

*M_k_* is parameterized by the mean (µ*_k_*) of the read arrival rate and the dispersion parameter (*R_k_*, related to the variance [σ^2^] of the read arrival rate by 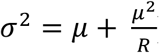, Γ(x) is the Gamma distribution and the read arrival rate (*λ_i_*) is defined as 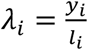 (*l_i_* is length of contig *x_i_*). To accommodate the presence of multiple contigs of different lengths in a component, we transform contigs into sets of windows of the same size (default=340bp) such that *Y* is replaced with the set of read counts for windows across all contigs from *X* (*Ω* = {ω_1_ … ω_u_}) in the model. Our goal then is to learn the mixture model *M* and the most probable assignment of contigs to the components of *M*, given the read counts *Y* and the dispersion parameter *R* (see below for how that is estimated).

In order to identify suitable mixture models we use the approach of computing Bayes factors to compare models as follows (where *Λ*_i_ are model parameters for component *M*_i_):

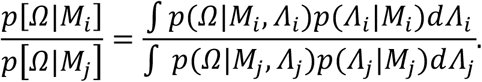

Defining a prior distribution for model parameters however requires information on the known relative abundance of genomes in the sample, which is typically not available in a *de novo* assembly setting. In addition, evaluation of the resulting multiple integrals, especially for the large number of potential models considered can be computationally intensive. To overcome these obstacles, we adopted the well-studied approach of using unit information priors (UIPs) as they allow for the approximation of the integrated likelihood using a closed form expression: 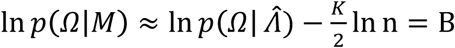, with 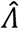 being the maximum-likelihood estimate (MLE) for the parameters, and *B* being a commonly used model selection score i.e. the *Bayesian Information Criterion* ^53,54^ (BIC). Spread-out priors like that of the *UIP* have the advantage of being conservative in that more evidence is needed from the data to support additional mixture components and hence components detected by the BIC are likely to be well supported by the data.

To constrain the selection of clusters, OPERA-MS first constructs a “guide tree” that encodes prior beliefs on possible partitions. It begins by assigning every contig *x_i_* to its own cluster *C_i_* with the average arrival rate of the cluster *λ_i_* set to the average for the contig (= *y_i_*/*l_i_*). Edges in the assembly graph are then weighted by the distance metric 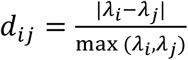, for adjacent clusters *C_i_* and *C_j_*, and then edges are traversed in increasing order of *d_ij_* to find contigs in different clusters that can potentially merge. In the resulting guide tree *T*, every internal node *n_k_* proposesa unique cluster *C_k_* (a mixture component) for all the contigs represented by its child nodes *c*(*n_k_*), with 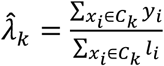 i.e. the maximum-likelihood estimate. The construction of the guide tree in this manner has several advantages: (i) firstly by relying on assembly edges, the guide tree eliminates models that cluster contigs from different connected components in the assembly graph; (ii) secondly, the distance between contigs in the guide tree increases with distances in coverage space as well as genomic location. Thus fewer internal nodes in the tree (and clustering models) propose clustering distant contigs together; (iii) finally, by considering edges in the order of the distance metric, potentially erroneous links are likely to be used closer to the root and thus are easier to eliminate in the model selection step as described below.

#### Cluster selection based on Bayesian Information Criterion

In order to compute the mixture model in *T* which maximizes the BIC, we exploit the fact that the BIC score is decomposable in a fashion that enables the use of a dynamic programming algorithm on a tree to optimize it (**Figure 1**). Specifically, 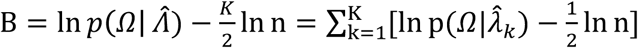 and hence the score of a *K* cluster decomposition of the tree can be expressed in terms of scores for each parent node. Thus adynamic programming algorithm that selects between the parent node and the children at each level of the tree can be used to efficiently find an optimal decomposition of the guide tree in *0*(*n*)time similar to the work by Navlakha *et al*^55^, based on the following recursion: 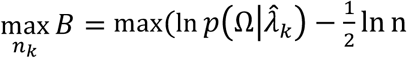, 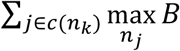

To select a suitable value for the dispersion parameter *R* for our models, we do a global search between the minimum estimate for a contig (lower-bounded at 1) and the joint estimate for all contigs larger than 10kbp (obtained via least squares regression), and using the smallest value where the corresponding clustering does not have a cluster larger than 200kbp for which 1% of the contig windows have coverage that varies two-fold compared to the mean (i.e. below *μ*_c_/2 or above 2*μ*_c_ when mean coverage is *μ*_c_). This search procedure allows us to use a global estimate for 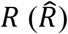, while reporting conservative clusters that do not merge multiple genomes together.

#### Computation of MASH genomic distance against a database of 2,800 complete genomes

The Bayesian model selection based approach for clustering is conservative and unsupervised by design, and for species where not enough sequencing data is present, can fragment the genome into multiple clusters. We therefore augment this with a supervised approach that compares all assembled sequences against a database of complete genomes (NCBI v2015.05.04) to merge clusters further. Specifically, we use the Mash toolkit^56^ based on the MinHash algorithm to efficiently estimate the similarity between contig clusters and reference genomes. The top 5 hits for each cluster with distance below 0.9 were then used to associate potential species names to them.

#### Identification of clusters from the same species

In order to super-cluster the original set of clusters, we first identified assembly edges between contigs that were supported by our Mash analysis. In particular, we required that when two contigs were aligned with MUMmer^57^ to the common reference genome identified by Mash, the estimated distance between them was within 3 standard deviations of the estimate provided by the assembly edge. This process was also allowed to rescue edges that had only 1 long read supporting them. All assembly graph edges were then used to merge clusters which had at least 1 species name common among their list of potential species names to construct species concordant super-clusters that had support in the assembly graph.

#### Sub-species level clustering

To account for species level clusters that potentially had multiple strain genomes merged, we used a further deconvolution step that worked on each such cluster separately. We first identified such clusters based on the total sequence size being >10% of the reference genome size of the associated species. Contigs belonging to different strains were then identified based on their read coverage as follows: i) the coverage distribution across contig windows as defined before was used to identify multi-modality based on a kernel density estimation approach to smooth the empirical distribution and infer local maxima^58^, and ii) for each mode *m* the mean coverage 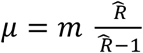 was computed and used to partition contigs based on a 96% confidence interval generated from a Negative Binomial distribution with parameters *μ* and 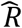 (preferentially assigning contigs to modes with higher coverage).

#### Pre- and post-processing

Unless otherwise stated, MegaHit contigs were used for further assembly and contigs shorter than 500bp were excluded. Estimation of contig coverage was performed by mapping short reads to the assembly using BWA-ALN^59^ (v0.7.10-r789, default parameters). Coverage was computed for 360bp windows, with the first and last 80bp of contigs being excluded to avoid edge effects. MASH (v1.1.1) was used to sketch the NCBI database of complete bacterial genomes (v2015.05.04). All mappings using MUMmer^57^ (v3.23) were performed with the parameter --max-match. Contig clusters were scaffolded and gap-filled using OPERA-LG (v1.0, with repeat detection disabled; species level clusters for mock community).

### Parameter settings for metagenomic assembly comparisons

Parameter settings for other benchmarked assemblers were as follows. MegaHit^21^ (v1.0.4-beta): default parameters; metaSPAdes^22^ (v3.7.1): -k 21,33,55,81; IDBA-UD^23^ (v1.1.1): default parameters; Canu^25^ (v1.5): the value of the parameter genomeSize was obtained by computing the MegaHit assembly size, -nanopore-raw for nanopore reads, -pacbio-raw otherwise; hybridSPAdes^26^ (v3.7.1): -k 21,33,55,81, --nanopore for nanopore reads, --pacbio otherwise.

Assemblies from MegaHit, metaSPAdes, IDBA-UD and OPERA-MS were polished using Pilon (v1.22, default parameters) with BWA-MEM (default parameters) mapping of Illumina reads to the assembly.Canu assemblies were further improved using the consensus module Racon^60^ (commit #0834442, default parameters) using GraphMap^61^ (v0.2.2, default parameters) mapping of the long reads to the assembly.

### Construction of GIS20 mock community

Six strains of bacteria were purchased from ATCC: *Pseudomonas putida* (ATCC^®^ 39213™), *Klebsiella pneumoniae* (ATCC^®^700721™), *Acinetobacter baumannii* (ATCC^®^ 17978™), S*taphylococcus epidermidis* (ATCC^®^ 12228™), *Enterococcus faecium* (ATCC^®^ BAA-472™) and *Salmonella enterica* (ATCC^®^ 13311™). Bacterial cells were cultured according to ATCC specified growth conditions. Overnight cultures were centrifuged at 4000*g* for 10min. Pelleted cells were stored at −20°C until DNA extraction was carried out. DNA was extracted using Genomic Tip 100/G (10243, Qiagen) and Genomic DNA Buffer Set (19060, Qiagen) according to the manufacturer’s protocol. DNA for the remaining 14 strains of bacteria was purchased from Leibniz Institute DSMZ (**Supplementary File 2**). Integrity of both extracted and purchased DNA was inspected on a 0.5% agarose gel and DNA concentration was determined using Qubit dsDNA BR assay (Q32853, Thermo Fisher Scientific). A staggered mock community was constructed by pooling DNA for the strains in a wide range of abundances varying from 0.1% to 30% (**Supplementary File 2**).

### Pacbio sequencing of GIS mock community

Library preparation for the mock community was performed according to “20kb Template Preparation Using BluePippin™ Size-Selection System”. Briefly, fragmentation of DNA was carried out using 20µg of DNA divided equally into 2 Covaris^®^ g-TUBE^®^ devices (520079, Covaris) in a total volume of 150µl each. The tubes were centrifuged with Eppendorf^®^ 5415R at 5,500 rpm for 1min and re-inserted in the opposite direction and centrifuged once more. The shearing process was repeated to obtain DNA fragments with sizes in the range 10-20kbp, yielding 16.2µg of DNA in total. Increased volume (3.24×) of reagents was used for downstream library preparation. The library was size-selected using the BluePippin™ Size-Selection System with a 0.75% DF Marker S1 high-pass 6-10kb cassette (BLF7510, Sage Science). The system protocol was set to select fragment sizes above 7kbp. The library profile was analyzed using Agilent DNA12000 Kit (5067-1508, Agilent Technologies). Eight SMRT cells were sequenced on the Pacbio RS II using P5 chemistry of run length 360.

### HMP mock community datasets

Illumina short reads^18^ (SRR2822454) and Illumina synthetic long reads^18^ (SRR2822457) were downloaded for the HMP staggered mock community from the SRA database (https://www.ncbi.nlm.nih.gov/sra). As PacBio sequencing data for the HMP staggered mock community was not available, we downloaded the PacBio HMP even mock community dataset (https://s3.amazonaws.com/datasets.pacb.com/Human microbiome mockB/hmp set5.tar.gz/hm p setf 5-7].tar.gz) and generated a staggered version using the following protocol: i) mapping reads to the reference genome using GraphMap (v0.2.2, default parameters), ii) computing for each genome the ratio *r_g_* of observed abundances *versus* expected abundances, iii) subsampling reads to obtain the same ratio *r_g_* for observed *versus* expected abundances in the staggered community.

### Evaluation of metagenomic assemblies

MetaQUAST^62^ (v 4.0, --genes) was used to evaluate metagenomic assemblies (in comparison to known reference genomes and annotations for mock communities) and obtain statistics such as NGA50 (aligned assembly length such that >50 % of the genome is in fragments of equal or greater length), genes assembled, misassembly errors and basepair errors.

Binning of metagenomic assemblies was performed using MaxBin2^10^ (v2.2.4, default parameters) and the quality of bins was assessed based on MUMmer mapping (from MetaQUAST output) to mock community reference genomes and calculation of the following metrics:

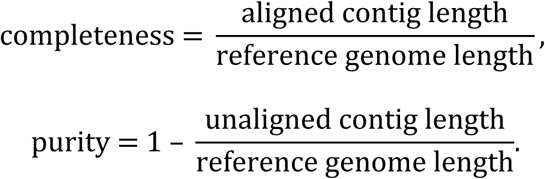

Binned assemblies were evaluated for their ability to recover complete biological pathways as follows: KAAS^63^ was used to identify KEGG orthology groups (KO) for genes in the reference genomes, and this annotation was used to establish the pathways present. MetaQUAST (v4.0, default parameters) was then used to identify genes present in the assembly using the --genes option, and these were then used to establish the fraction of genes assembled per pathway and the fraction of pathways that are captured well in the binned assemblies.

### Analysis of mobile elements and resistance genes

Plasmid and phage sequences that were likely fully assembled were identified by looking for evidence for large circular sequences in the assembly. Specifically, we mapped the long reads back to the OPERA-MS assembly using BLASR^50^ (v1.3.1, -m 1 --minMatch 5 --bestn 10) and checked for the presence of at least 2 reads that spanned contig ends (reads mapped <400bp away from contig ends). In addition, putative plasmids and phage sequences were then mapped against the nt database using blastn^64^ (2.2.28+, default parameters) and sequences with chained best blast hits aligning more than 50% of the sequence to non plasmid/phage hits were excluded as potential misassemblies. Partially assembled plasmid sequences were identified based on contigs that were >20kbp long where Mash and blastn identified a plasmid sequence as their best hit covering >90% of their sequence. Antibiotic resistance (AR) genes were identified by aligning sequences >5kbp to the ARG-ANNOT^65^ database (V2) using blastn. AR genes with >98% of the sequence aligning to the contig with an identity >99% were selected for further analysis.

### Annotation and phylogenetic analysis for the putative jumbo phage

The assembled sequence was analyzed using the RAST server^66^ to identify and annotate putative genes. Sequences for the DNA polymerase B protein from bacteria, small bacteriophages and jumbo phages were aligned with MUSCLE^67^ (v3.8.31, default parameters) and a phylogenetic tree (500 bootstrap replicates) was generated using PhyML^68^ and the pipeline provided at http://www.phylogeny.fr/.

### Analysis of strain dynamics

To estimate the abundance of the two *K. pneumoniae* strains that were obtained from OPERALG’s strain level assembly, Illumina reads were mapped back to the assemblies with stringent criteria (BWA-MEM, mappings with soft clipping or >3 mismatches filtered). The strains with higher and lower coverages at the time-point for which the hybrid assembly was constructed were denoted as strain H and L respectively. The median coverage of 500bp windows in the two assemblies was then computed (*c_H_*, *c_L_*), and the abundance of strain L was estimated as *c_L_* while the abundance of strain H was estimated as *c_H_* − *c_L_* (to account for the fact that the majority of shared contigs for the strains will be incorporated into the strain H assembly).

### Confirmation of plasmid association

CRE isolates were obtained from stool DNA as described before^69^ and Genomic DNA was extracted from an overnight culture using MagNA Pure Compact (Roche Applied Science, Germany). Library preparation was performed using the NEBNext Ultra™ DNA Library Prep Kit and 2×151bp sequencing was performed using the Illumina HiSeq 4000. *De novo* assembly was performed using Velvet^70^ (version 1.2. 10) with parameters optimized by Velvet Optimiser (k-mer ranging from 81 to 127), scaffolded with OPERA-LG^24^ (version 1.4.1), and finished with FinIS^71^ (version 0.3). MUMmer^57^ alignments were used to confirm the presence of the plasmid, high identity to the genome of strain L (>99.9%) and low identity to the genome of strain H (<97.5%).

### Availability of data and software

GIS20 mock community sequencing data can be obtained from the European Nucleotide Archive (ENA) under project ID PRJEB29139 (Illumina, PacBio and ONT) and sequencing data for the 28 gut metagenomes can be found under project ID PRJEB29152 (Illumina and ONT). OPERA-MS is freely available under the MIT license at https://github.com/CSB5/OPERA-MS.

